# Characterization of Dyspnea in Lung Cancer Survivors Following Curative-Intent Therapy

**DOI:** 10.1101/508440

**Authors:** Duc Ha, Andrew L. Ries

**Affiliations:** Institute for Health Research; Kaiser Permanente Colorado; Aurora, Colorado; Division of Pulmonary, Critical Care, and Sleep Medicine; Department of Medicine; University of California, San Diego, La Jolla, California

**Keywords:** Lung neoplasms, patient-reported outcome measures, symptom assessment, exercise, survivorship

## Abstract

**Purpose:** Dyspnea is highly-prevalent in lung cancer survivors following curative-intent therapy. We aimed to identify clinical determinants of dyspnea and characterize its relationship with functional exercise capacity (EC).

**Methods:** In an analysis of data from a cross-sectional study of lung cancer survivors who completed curative-intent therapy for stage I-IIIA disease ≥1 month previously, we tested a thorough list of comorbidities, lung function, and lung cancer characteristics. We assessed dyspnea using the European Organization for Research and Treatment of Cancer Quality of Life Questionnaire Lung Cancer Module 13 (LC13) and functional EC the six-minute walk. We verified results with the University of California San Diego Shortness of Breath Questionnaire (SOBQ).

**Results:** In 75 participants at a median of 12 months since completing treatment, the mean (SD) LC13-Dyspnea score was 35.3 (26.2); 60% had abnormally-high dyspnea. In multivariable linear regression analyses, significant clinical determinants of dyspnea were [β (95% confidence interval)]: psychiatric illness [−20.8 (−32.4, −9.09) for *No/Yes*], heart failure with reduced ejection fraction [−15.5 (−28.0, −2.97) for *No/Yes*], and forced expiratory volume in 1 second [−0.28 (−0.49, −0.06) for each *% predicted*]. Dyspnea was an independent predictor of functional EC [−1.54 (−2.43, −0.64) for each *point*]. These results were similar with the SOBQ.

**Conclusion:** We identified clinical determinants of dyspnea which have pathophysiological bases. Dyspnea was independently associated with functional EC. Behavioral interventions to promote exercise in lung cancer survivors following curative-intent therapy may need to also optimize medical therapy for cardiopulmonary and/or psychiatric disease and reduce dyspnea to be effective.

## Introduction

Physical exercise has been shown to be effective in improving function and QoL in cancer survivors [1] and is recommended by the American College of Sports Medicine [2] and American Cancer Society [3] for cancer survivors. However, the evidence of effectiveness is not as consistent in lung cancer survivors [4]. This inconsistency may be related to characteristics unique to lung compared to other cancer survivors, including differences in age, comorbidities, and the effects of lung cancer and its treatment on exercise capacity (EC). Many lung cancer patients are elderly and have major comorbidities that include chronic obstructive pulmonary disease (COPD) and heart failure (HF) which may limit EC [5]. Also, curative-intent therapy of lung cancer necessitates the removal and/or destruction of lung tissue which may further impede cardiopulmonary function and EC in some patients.

Symptom burden may limit physical exercise in lung cancer survivors. Following lung cancer resection surgery, dyspnea worsens [6] partly due to a loss of 10-15% of lung function [7]. Clinically significant dyspnea has been shown to be prevalent in >50% of lung cancer survivors [8–11]. As a result, lung cancer survivors may avoid physical exercise to prevent dyspnea. In time, this avoidance can lead to a downward spiral of health related to physical function, symptom burden, and QoL (E-Figure 1), all of which are important survivorship issues in lung cancer [12]. Therefore, the characterization of dyspnea in lung cancer survivors may provide important insights into factors associated with dyspnea and may facilitate interventions to improve exercise, function, and QoL in these patients.

**Figure 1:**
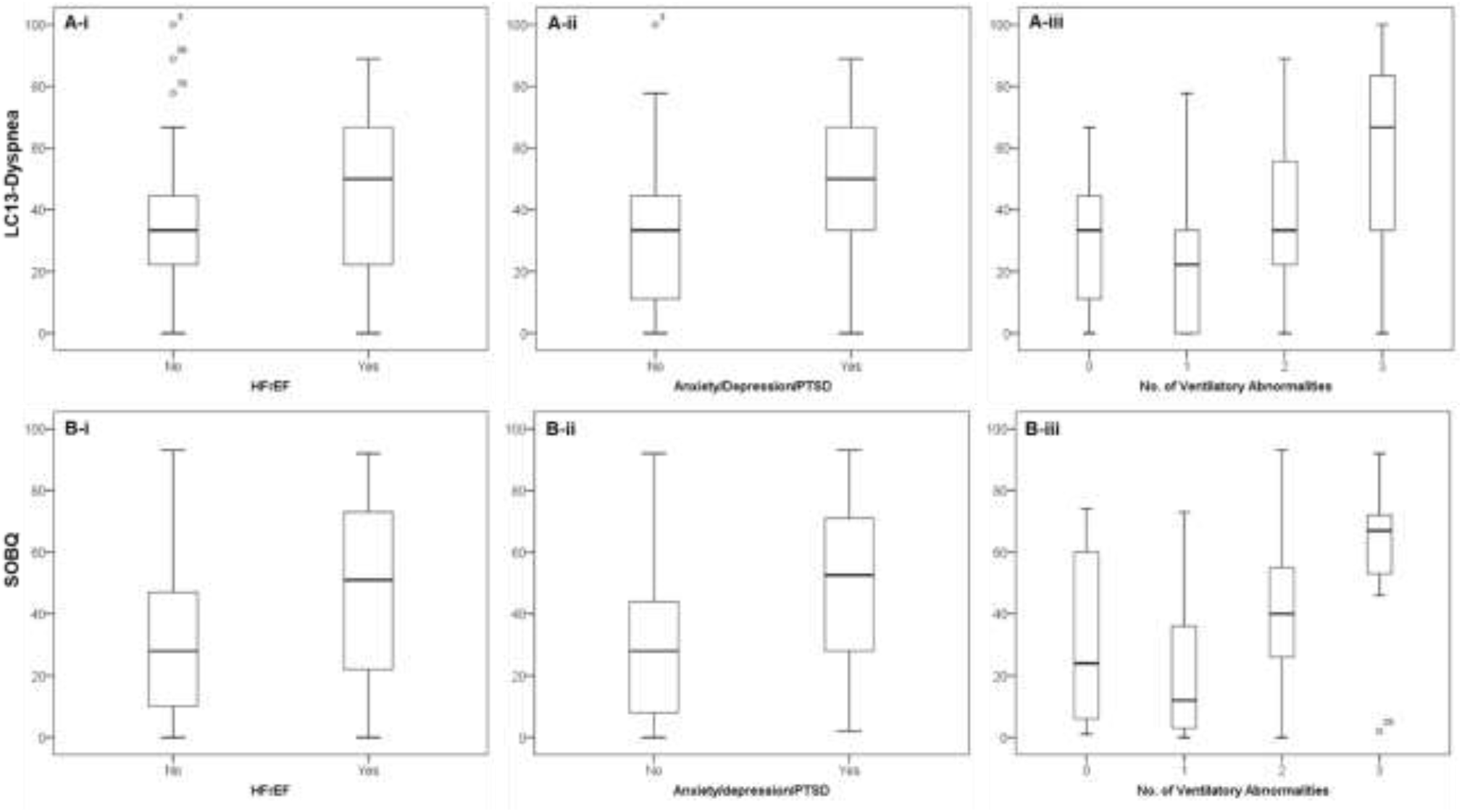
Clinical Determinants of Dyspnea. (A) Dyspnea scores as assessed by the LC13, stratified by: 
i. HFrEF; mean difference (standard error, SE) = 15.3 (6.91), p* = 0.03.
ii. Anxiety/depression/PTSD; mean difference (SE) = 19.4 (6.30), p* = 0.003.
iii. Combined ventilatory abnormalities; one-way analysis of variance (ANOVA) F-statistics = 3.22, p = 0.03; statistically significant difference between 3 compared to 1 ventilatory abnormalities [(mean difference (SE) = +32.3 (11.0), p** = 0.03]. (B) Dyspnea scores as assessed by the SOBQ, stratified by: 
i. HFrEF; mean difference (SE) = 16.5 (6.69), p* = 0.02.
ii. Anxiety/depression/PTSD; mean difference (SE) = 19.9 (6.10), p* = 0.002.
iii. Combined ventilatory abnormalities; one-way ANOVA F-statistics = 6.03, p = 0.001; statistically significant difference between 3 compared to 1 [mean difference (SE) = +39.0 (10.2), p** = 0.002] and 2 compared to 1 ventilatory abnormalities [mean difference (SE) = +20.0 (6.4), p** = 0.01]. * *Independent sample t-tests, equal variances assumed.* ** *Bonferroni-corrected for multiple comparisons*. ANOVA = one-way analysis of variance; LC13 = European Organization for Research and Treatment of Cancer QoL Questionnaire Lung Cancer Module 13; HFrEF = heart failure with reduced ejection fraction; PTSD = post-traumatic stress disorder; SE = standard error; SOBQ = University of California San Diego Shortness of Breath Questionnaire

In this project, we aimed to identify clinical determinants of dyspnea in lung cancer survivors following curative-intent therapy and analyze its relationship with functional EC. We believe that optimization of underlying comorbidities including cardiopulmonary disease may be important in improving physical exercise, function, and QoL in these patients.

## Methods

### Study Overview

We previously performed an Institutional Review Board-approved, crosssectional study to assess functional EC and patient-reported outcomes (PROs) in lung cancer survivors following curative-intent therapy [13]. In exploratory PRO assessments, we identified abnormally-high dyspnea in approximately 60% of participants [13] as assessed by the European Organization for Research and Treatment of Cancer QoL Questionnaire Lung Cancer Module 13 (LC13) [14]. In this study, we enrolled additional participants to identify clinical determinants of dyspnea and explore its relationship with functional EC. We followed guideline recommendations for Strengthening the Reporting of Observational studies in Epidemiology to report our findings [15].

### Participants & Variables

We previously reported our enrollment methods, inclusion, and exclusion criteria, through which we enrolled ~87% of all eligible patients [13]. In this study, we excluded patients with severe dementia (n=2), bilateral below-knee amputation (n=2), quadriplegia (n=1), or active systemic treatment for other cancers (n=4). We abstracted clinical variables related to lung cancer and/or cardiopulmonary health. These included: age, sex, body-mass index, tobacco exposure, comorbidities (including COPD, HF, psychiatric illness), lung function [forced expiratory volume in 1 second (FEV_1_), diffusion of the lung for carbon monoxide (DL_CO_), total lung capacity (TLC)], and echocardiographic findings where available. Lung cancer characteristics included clinical stage, treatment modality, and time since treatment completion.

### Dyspnea Assessments

We assessed dyspnea using the LC13 [14] and the University of California San Diego Shortness of Breath Questionnaire (SOBQ) [16]. We used the LC13-Dyspnea score for primary analyses due to previous validation in the lung cancer population [14] and the SOBQ to verify our findings. The LC13 is a 13-item questionnaire designed to assess lung cancer-associated symptoms (cough, dyspnea, hemoptysis, and pain) and chemo- and radiotherapy side-effects (dysphagia, hair loss, neuropathy, sore mouth); dyspnea is assessed by three items on perceived shortness of breath at rest, when walking, and climbing stairs, scored as a mean of the component items with raw score range 0-100 [17]. We defined abnormally-high dyspnea as LC13-Dyspnea scores >mean reference value [18]. The SOBQ is a 24-item questionnaire designed to assess self-reported shortness of breath with a variety of activities of daily living in patients with chronic lung disease [16]; responses are summed to a score range 0-120 [16], In both the L13 and SOBQ questionnaires, higher scores indicate a higher level of perceived dyspnea. We performed all questionnaire assessments at the same time on printed forms without any modifications.

### Functional EC

We defined our primary outcome as functional EC as assessed by the distance covered (6MWD) during the six-minute walk test (6MWT). In lung cancer survivors, the 6MWD has been validated against the gold standard of cardiopulmonary fitness [19] shown to decrease with treatment [20], low compared to general age-, sex-, height-, and weight-matched adults [13], and independently associated with cancer-specific QoL [13]. We performed the 6MWT according to the standard protocol at our institution which follows the American Thoracic Society (ATS) Pulmonary Function Standards Committee recommendations [21].

### Statistical Analyses

We analyzed both L13-Dyspnea and SOBQ scores as continuous variables. We tested all collected clinical characteristics to identify determinants of dyspnea and analyzed the relationship between dyspnea and functional EC. We also assessed an exploratory physiological determinant of combined ventilatory abnormalities, defined as any combination of obstructive ventilatory defect, lung hyperinflation, and DL_CO_ limitation on dyspnea.

We summarized descriptive statistics as appropriate. We used univariable (UVA) and multivariable (MVA) linear regression analyses to identify clinical determinants of dyspnea scores and predictors of functional EC; for MVAs, we used stepwise, backward selection modeling starting with all variables with p <0.10 in UVAs except where indicated. We interpreted results using regression coefficients (β), 95% confidence intervals (CIs), and coefficients of determination (R^2^ and partial R^2^), and defined statistical significance as p <0.05 in two-tailed tests. We corrected p-values for multiple comparisons where applicable. All data were managed using REDCap electronic data capture tools hosted at the UCSD Clinical and Translational Research Institute [22] and analyzed by IBM^®^ SPSS^®^ Statistics software version 25.

## Results

### Participants

We enrolled 75 lung cancer survivors at a median of 12 months since completing curative-intent therapy. Their baseline clinical characteristics as described in Table 1. Most were white males with a history of tobacco use, concomitant COPD, and underwent either lung cancer resection surgery or definitive radio-ablation for treatment of stage I non-small cell lung cancer (NSCLC).

**Table 1:** Participant Characteristics

| Participant Characteristic (N=75) | Value |
| --- | --- |
| Age, mean (SD) | 70.7 (8.4) |
| Male sex, n (%) | 71 (95) |
| Race, n (%) |  |
| Asian | 4 (5) |
| Black | 2 (3) |
| Hispanic | 1 (1) |
| White | 68 (91) |
| BMI, kg/m<sup>2</sup>, mean (SD) | 26.8 (4.8) |
| Smoking history, n (%) |  |
| Current | 24 (32) |
| Former | 44 (59) |
| Never | 7 (9) |
| Pack years, mean (SD) | 52.0 (31.7) |
| Comorbidities, n (%) |  |
| Hypertension | 61 (81) |
| Hyperlipidemia | 60 (80) |
| Diabetes mellitus | 21 (28) |
| Atrial fibrillation/flutter | 18 (24) |
| CAD | 26 (35) |
| COPD | 56 (75) |
| HFrEF<sup>†</sup> | 18 (24) |
| Diastolic dysfunction<sup>*</sup> | 37 (79) |
| OSA | 17 (23) |
| Anxiety/depression/PTSD | 22 (29) |
| Other cancer | 30 (40) |
| Pulmonary function<sup>**</sup>, mean (SD) |  |
| FEV<sub>1</sub>/FVC % | 59.6 (14.6) |
| FEV<sub>1</sub> % predicted | 70.7 (25.0) |
| TLC % predicted<sup>*</sup> | 110.8 (21.5) |
| DL<sub>CO</sub> % predicted | 78.5 (25.4) |
| Ventilatory abnormality<sup>‡</sup>, n (%) |  |
| Obstructive | 57 (76) |
| Hyperinflated | 18 (28) |
| DL<sub>CO</sub> limited | 41 (55) |
| Lung cancer |  |
| Clinical stage, n (%) |  |
| IA | 50 (67) |
| IB | 9 (12) |
| IIA | 3 (4) |
| IIB | 11 (15) |
| IIIA | 2 (3) |
| Histology, n (%) |  |
| Adenocarcinoma | 37 (49) |
| Squamous cell carcinoma | 18 (24) |
| Presumed | 15 (20) |
| Primary treatment, n (%) |  |
| Surgical resection | 37 (49) |
| Definitive radio-ablation | 27 (36) |
| Chemoradiation | 11 (15) |
| Months since treatment |  |
| Median | 12.2 |
| Interquartile range | 1.8 – 35.6 |
| Dyspnea scores, mean (SD) |  |
| LC13-Dyspnea, point | 35.3 (26.2) |
| SOBQ, point | 34.8 (25.6) |
| Functional EC (6MWD) *m*, mean (SD) | 347.9 (124.2) |

\*Missing data; data available in 64 participants (85%) for TLC % predicted and 47 (63%) for diastolic dysfunction.
\*\*Data obtained after completion of lung cancer treatment in 32 (43%) participants.
<sup>†</sup>Defined as ejection fraction $\leq 50\%$ or clinical documentation of systolic heart failure.
<sup>‡</sup>Defined as $FEV_1/FVC < 70\%$ for obstructive defect, TLC % predicted $> 120$ for hyperinflation, and $DL_{CO}$ % predicted $< 80$ for $DL_{CO}$ limitation.
6MWD = six-minute walk distance; LC13 = European Organization for Research and Treatment of Cancer QoL Questionnaire Lung Cancer Module 13; BMI = body-mass index; CAD = coronary artery disease; COPD = chronic obstructive pulmonary disease; $DL_{CO}$ = diffusion capacity of the lung for carbon monoxide; EC = exercise capacity; $FEV_1$ = forced expiratory volume in 1 second; FVC = forced vital capacity; HFrEF = heart failure with reduced ejection fraction; OSA = obstructive sleep apnea; PTSD = post-traumatic stress disorder; QoL = quality of life; SD = standard deviation; TLC = total lung capacity; SOBQ = University of California San Diego Shortness of Breath Questionnaire

### Dyspnea and Functional EC

All participants completed dyspnea and 6MWT assessments. The mean’s (SD’s) LC13-Dyspnea and SOBQ scores were 35.3 (26.2) and 34.8 (25.6), respectively. These scores correlated very well (r = 0.84, p <0.001) (E-Figure 2). Forty-five participants (60%) had abnormally-high dyspnea, defined as LC13-Dyspnea raw score >mean reference value for lung cancer patients (29.5) [18]. The mean (SD) 6MWD was 347.9 (124.2) m (67% of predicted values in healthy adults [23]) (Table 1).

**Figure 2:**
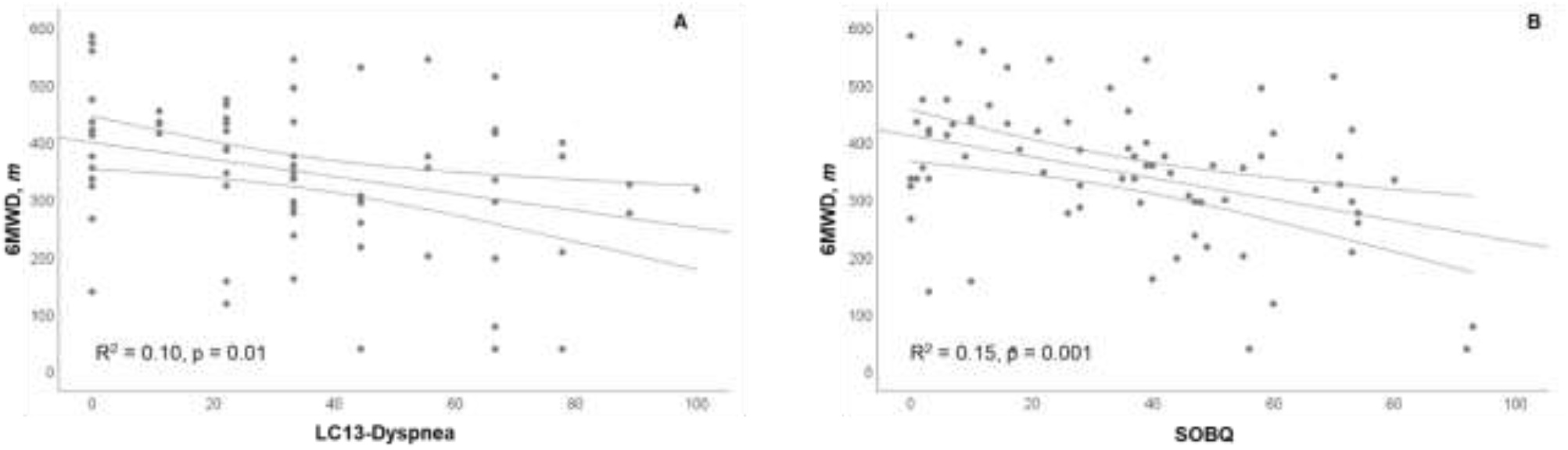
Dyspnea is an Independent Predictor of Functional EC (6MWD) (A) Dyspnea as assessed by the LC13 (B) Dyspnea as assessed by the SOBQ *R*^2^ *values derived from UVAs as listed in E-Table 2* 6MWD = six-minute walk distance; EC = exercise capacity; LC13 = European Organization for Research and Treatment of Cancer QoL Questionnaire Lung Cancer Module 13; QoL = quality of life; SOBQ = University of California San Diego Shortness of Breath Questionnaire; UVA = univariable linear regression analysis

### Determinants of Dyspnea

Clinical variables borderline-significantly associated with dyspnea are shown in E-Table 1. In MVAs [β (95% CI], psychiatric illness [−20.8 (−32.4, −9.09) for *No/Yes*], heart failure with reduced ejection fraction (HFrEF) [−15.5 (−28.0, −2.97) for *No/Yes*], and FEV_1_ [−0.28 (−0.49, −0.06) for each *% predicted*] were significant independent determinants of dyspnea. Results were similar with the SOBQ (Table 2A, Figure 1A&B-i-ii).

**Table 2:** MVA – Determinants of Dyspnea

(A) Independent Determinants of Dyspnea
| *LC13-Dyspnea<sup>†</sup>* |  |  |  |
| --- | --- | --- | --- |
| Variable | $\beta$ (95% CI) | Partial R<sup>2</sup> | P value |
| HFrEF (N/Y) | -15.5 (-28.0, -2.97) | 0.08 | 0.02 |
| Anxiety/depression/PTSD (N/Y) | -20.8 (-32.4, -9.09) | 0.15 | 0.001 |
| FEV<sub>1</sub> % predicted | -0.28 (-0.49, -0.06) | 0.09 | 0.01 |
| *SOBQ<sup>‡</sup>* |  |  |  |
| HFrEF (N/Y) | -15.8 (-27.9, -3.63) | 0.10 | 0.01 |
| Anxiety/depression/PTSD (N/Y) | -17.0 (-29.0, -4.91) | 0.12 | 0.01 |
| Lung hyperinflation* (N/Y) | -11.0 (-23.3, 1.21) | 0.05 | 0.08 |
| FEV<sub>1</sub> % predicted | -0.34 (-0.56, -0.11) | 0.13 | 0.004 |

\*Data available in 64 participants.
<sup>†</sup>Overall model R<sup>2</sup> = 0.26, $p < 0.001$ ; no significant interaction between HFrEF and anxiety/depression/PTSD ( $p = 0.14$ ) or anxiety/depression/PTSD and FEV<sub>1</sub> % predicted ( $p = 0.48$ ).
<sup>‡</sup>Overall model R<sup>2</sup> = 0.37, $p < 0.001$ ; no significant interaction between HFrEF and anxiety/depression/PTSD ( $p = 0.15$ ), or anxiety/depression/PTSD and FEV<sub>1</sub> % predicted ( $p = 0.68$ ).

(B) Combined Ventilatory Abnormalities as an Exploratory Determinant of Dyspnea
| *LC13-Dyspnea<sup>†</sup>* |  |  |  |
| --- | --- | --- | --- |
| Variable | $\beta$ (95% CI) | Partial R<sup>2</sup> | P value |
| HFrEF (N/Y) | -15.0 (-28.2, -1.79) | 0.07 | 0.03 |
| Anxiety/depression/PTSD (N/Y) | -17.5 (-30.4, -4.62) | 0.10 | 0.01 |
| Combined ventilatory abnormality* | N/A | 0.06 | 0.23 |
| *SOBQ<sup>‡</sup>* |  |  |  |
| HFrEF (N/Y) | -15.2 (-27.3, -2.99) | 0.08 | 0.02 |
| Anxiety/depression/PTSD (N/Y) | -15.8 (-27.7, -3.83) | 0.09 | 0.01 |
| Combined ventilatory abnormality* | N/A | 0.12 | 0.03 |

\*Defined as any combination of obstructive ventilatory defect, lung hyperinflation, and DL<sub>CO</sub> limitation; range 0-3.
<sup>†</sup>Overall model R<sup>2</sup> = 0.24, $p = 0.001$ .
<sup>‡</sup>Overall model R<sup>2</sup> = 0.32, $p < 0.001$ .
DL<sub>CO</sub> = diffusion capacity of the lung for carbon monoxide; LC13 = European Organization for Research and Treatment of Cancer QoL Questionnaire Lung Cancer Module 13; FEV<sub>1</sub> = forced expiratory volume in 1 second; HFrEF = heart failure with reduced ejection fraction; PTSD = post-traumatic stress disorder; SOBQ = University of California San Diego Shortness of Breath Questionnaire

In exploratory UVAs, combined ventilatory abnormalities of obstructive ventilatory defect, lung hyperinflation, and DL_CO_ limitation (categorized as a range of 0-3) was a significant determinant of dyspnea in both LC13-Dyspnea (p = 0.03) and SOBQ (p = 0.001) scores. Post-hoc analyses with Bonferroni correction for multiple comparisons showed that those with 1 and 2 ventilatory abnormalities had significantly lower dyspnea scores than those with 3 ventilatory abnormalities (Figure 1A&B-iii, mean differences provided in figure legend). Adjusting for HFrEF and psychiatric illness, combined ventilatory abnormality was a significant independent determinant of dyspnea as assessed by the SOBQ (partial R^2^ = 0.12, p = 0.03), but not LC13-Dyspnea in MVAs (Table 2B).

### Dyspnea is an Independent Predictor of Functional EC

Results of UVAs to identify predictors of functional EC are shown in E-Table 2. In MVAs starting with all baseline clinical characteristics borderline-significantly associated with functional EC, dyspnea was found to be an independent predictor of functional EC [−1.54 (−2.43, −0.64) for each *point]* (Table 3) (Figure 2A). These results were replicated by the SOBQ [−1.70 (−2.62, −0.78) (Table 3) (Figure 2B).

**Table 3:** Dyspnea is an Independent Predictor of Functional EC

| *Model 1<sup>†</sup>* |  |  |  |
| --- | --- | --- | --- |
| Variable | $\beta$ (95% CI) | Partial R<sup>2</sup> | P value |
| Age, year | -5.81 (-8.54, -3.08) | 0.21 | <0.001 |
| Hyperlipidemia (N/Y) | 72.0 (15.9, 128.0) | 0.09 | 0.01 |
| DL<sub>CO</sub> % predicted | 1.81 (0.90, 2.73) | 0.18 | <0.001 |
| LC13-Dyspnea, point | -1.54 (-2.43, -0.64) | 0.14 | 0.001 |
| *Model 2<sup>‡</sup>* |  |  |  |
| Age, year | -5.86 (-8.55, -3.16) | 0.21 | <0.001 |
| Hyperlipidemia (N/Y) | 61.6 (6.62, 116.6) | 0.07 | 0.03 |
| DL<sub>CO</sub> % predicted | 1.70 (0.79, 2.62) | 0.16 | <0.001 |
| SOBQ, point | -1.70 (-2.62, -0.78) | 0.16 | <0.001 |

<sup>†</sup>Overall model R<sup>2</sup> = 0.45, $p < 0.001$ ; no significant interaction between age and LC13-Dyspnea ( $p = 0.40$ ), or DL<sub>CO</sub> % predicted and LC13-Dyspnea ( $p = 0.30$ ).
<sup>‡</sup>Overall model R<sup>2</sup> = 0.46, $p < 0.001$ ; no significant interaction between age and SOBQ ( $p = 0.995$ ), or DL<sub>CO</sub> % predicted and SOBQ ( $p = 0.55$ ).
<sup>†‡</sup>Covariates selected to enter model ( $p < 0.1$ ): age, hyperlipidemia, CAD, FEV<sub>1</sub> % predicted, DL<sub>CO</sub> % predicted, adenocarcinoma, treatment group (E-Table 3).
CAD = coronary artery disease; CI = confidence interval; DL<sub>CO</sub> = diffusion capacity of the lung for carbon monoxide; EC = exercise capacity; LC13 = European Organization for Research and Treatment of Cancer QoL Questionnaire Lung Cancer Module 13; FEV<sub>1</sub> = forced expiratory volume in 1 second; SOBQ = University of California San Diego Shortness of Breath Questionnaire

## Discussion

In an analysis of data from a cross-sectional study of veteran lung cancer survivors following curative-intent therapy, we found abnormally-high dyspnea in most patients and identified three important clinical determinants of dyspnea: psychiatric illness (anxiety/depression/PTSD), HFrEF, and physiological measure of obstructive ventilatory defect (FEV_1_). In addition, dyspnea was an independent predictor of functional EC and accounted for 14-16% of its variance. These results highlight the importance of dyspnea in lung cancer survivorship and may have implications for interventions aimed at improving physical exercise, function, and QoL in these patients.

Exercise training is recommended by the American College of Sports Medicine as an adjunct cancer therapy [2] and regular physical activity is recommended by the American Cancer Society [3] for cancer survivors. These recommendations are based on evidence of effectiveness of physical exercise in improving symptom control, function, and QoL, mostly in breast, prostate, and colon cancer survivors. The evidence for effectiveness is not as consistent in lung cancer survivors [4]. Recent systematic reviews of exercise interventions in lung cancer survivors suggest that while exercise training can improve functional EC and QoL following curative-intent therapy [24–27], there are concerns for bias, inadequate sample size, and lack of large randomized controlled trials [24, 25, 28]. Moreover, a recent systematic review identified numerous patient-related barriers to regular physical activity in lung cancer survivors [29]. These include physical capacity, symptoms, comorbidities, previous sedentary lifestyle, and psychological influences [29].

Lung cancer survivors are different from other cancer populations due to a higher median age at diagnosis, lifetime exposure to tobacco, and prevalence of pulmonary and cardiovascular comorbidities [5]. The median age at lung cancer diagnosis is 70 years [30]. Eighty to 90% lung cancer cases are caused by cigarette smoking [31] which has many adverse health consequences. The most common comorbidities in patients with lung cancer – all of which can progressively and negatively impact health – are chronic obstructive pulmonary disease (COPD, in 50-70% of patients [32, 33]), diabetes (16%), and HF (13%) [5]. In addition, the effects of curative-intent therapy of lung cancer have unique effects on cardiopulmonary health since part of the therapy requires local destruction and/or removal of lung tissue which may impede function and EC in some patients. Therefore, lung cancer survivors face unique health challenges that may make typical exercise interventions inaccessible [27]. The characterization of dyspnea may provide important insights into effective strategies to improve physical exercise in this group of cancer survivors.

The mechanisms of dyspnea are incompletely understood [34]; however, peripheral sensors including through reflex chemoreceptor stimulation by carbon dioxide, pulmonary vagal C-fibers, mechanoreceptors, and central pathways (specifically the limbic system and sensorimotor cortex) are thought to play important roles [34]. In lung cancer survivors following curative-intent therapy, dyspnea can be made worse due to a loss of 10-15% of lung function from lobectomy [35]. Additionally, other possible pathophysiological effects may include a loss of associated nerve fibers and peripheral sensors due to the removal and/or destruction of lung tissue by surgery and/or radiotherapy. Adjuvant chemotherapy has been shown to cause peripheral neuropathy [36] and, therefore, possibly also vagal perturbations. These effects may accumulatively result in neuromechanical dissociation which is implicated in the pathogenesis of dyspnea [37].

Importantly, dyspnea was independently associated with functional EC in our study. For each 1-point increase in baseline dyspnea score, there was a 1.5 – 1.7 *m* decrease in the 6MWD. The association between dyspnea and functional EC has been described in other patient populations including COPD [38]. To the best of our knowledge, this relationship has not been previously characterized in lung cancer survivors following curative-intent therapy, many of whom experience worsening of dyspnea [6] as laid out above. Moreover, a previous analysis of 359 post-surgical, stage I NSCLC survivors showed that dyspnea was significantly associated with physical health [10]. In fact, of all the variables included in this analysis, dyspnea had approximately twice the effect size (β) on physical health as the next two largest effect sizes (employment status and depression symptoms) [10]. Another study also confirmed this large effect size of dyspnea on physical health [39]. These findings suggest that alleviating dyspnea may be a key factor to improve physical exercise in these patients.

The prevalence of abnormally high dyspnea in our study (60%) is similar to previous reports in lung cancer survivors following curative-intent therapy [8–11]. In one of these studies [8], the prevalence of COPD (23%) was significantly lower than ours (75%), while the prevalence of abnormal/clinically significant dyspnea was the same. In that study, 65% dyspneic patients reported not having dyspnea preoperatively, suggesting treatment-related effects [8]. The determinants of dyspnea we identified offer complimentary information to a previous report in 342 post-surgical NSCLC survivors (preoperative dyspnea, preoperative DL_CO_, lack of moderate or strenuous physical activity, depression symptoms) [8], and another in 142 longterm, post-surgical NSCLC survivors (number of comorbidities, presence of moderate-severe ventilatory abnormality) [39]. We additionally identified HFrEf and psychiatric illness as determinants, which have clinical management implications in these patients. Also as in our study, clinical and physiological variables explained only approximately 30% of the variances in dyspnea scores [8],suggesting alternative etiologies not assessed (e.g. cardiopulmonary reserve [37]). Our exploratory analysis suggested that combined ventilatory abnormalities related to COPD (obstructive ventilatory defect, lung hyperinflation, and DL_CO_ limitation) may have additive negative impact on dyspnea, similar to a previous report in long-term survivors [39].

Moreover, LC13 dyspnea scores were recently reported to be higher in 830 post-surgical lung cancer survivors compared to 1,000 propensity-matched individuals from a Korean general population [40]. As well, adjusted dyspnea scores were higher in those treated with combined-modality therapy compared to those treated by surgery alone, and in those with cardiac or pulmonary comorbidity compared to those without comorbidity [40]. While our results showed an association between treatment type and dyspnea scores only in univariable but not multivariable analyses, possibly due to a small sample size and/or inclusion of predicted lung function in our analyses, the results laid out above suggest a complex relationship between dyspnea, comorbidities, and physical exercise behavior.

Based on a postulate of downward spiral of health resulting from curative-intent therapy (E-Figure 1), efforts to improve physical exercise, function, and QoL in lung cancer survivors may need to first optimize medical therapy for cardiopulmonary disease [41] and possibly psychiatric illness [42] to alleviate dyspnea to be effective. Recent randomized controlled trials in COPD patients show that combination bronchodilator inhaler therapy is effective in reducing dyspnea [43] and possibly in improving physical activity [44, 45]. Initiation of these inhalers, particularly in survivors with concomitant COPD, may be an important first step.

Our study has limitations. First, despite a thorough list of important clinical characteristics in the lung cancer population, we were only able to explain 30-40% of the variances in dyspnea scores. However, to the best of our knowledge, no other studies have reported a higher R^2^ values including those reported in a large, international, population based study involving 9,484 participants with many demographic and clinical variables (largest R^2^ reported = 0.13) [46]. Second, the cross-sectional and descriptive nature of our study provides no insight on the effects of lung cancer treatment on dyspnea, the underlying physiobiological mechanism, or how to effectively alleviate it. Third, our findings may have limited generalizability due to it being a single-institutional study involving a predominantly white male veteran patient population with significant tobacco exposure and higher prevalence of COPD and psychiatric illness [5]. Fourth, lung function (FEV_1_) but not a spirometric diagnosis of COPD was associated with dyspnea in our study. This may be due to our small sample size and/or a limitation in our data abstraction which contained pre- and post-treatment lung function tests.

The strengths of our study include a detailed list of comorbidities and physiological measures of cardiopulmonary health including lung function, all of which were entered in the electronic medical record and collected by board-certified physicians, maximizing the accuracy of the data obtained. In addition, all PRO and functional EC assessments were performed inperson by one observer (DH), maximizing the completeness and accuracy of the data collected and minimizing inter-observer variability. Last, we verified our findings using a second dyspnea questionnaire, maximizing the validity of our conclusions.

We conclude that abnormally-high dyspnea was prevalent in at least 50% of lung cancer survivors eligible following curative-intent therapy, partly due to underlying psychiatric illness, presence of HF, and obstructive ventilatory defects. In addition, dyspnea was a significant independent predictor of functional EC. Efforts to improve exercise, function, and QoL in lung cancer survivors may need to also focus on optimizing therapy for comorbid cardiopulmonary and/or psychiatric illnesses and reduce dyspnea to be effective.

## Disclosures and Acknowledgements

### Conflict of Interest

All authors declare no conflict of interest exists.

### Funding

This work was supported directly by the American Cancer Society (PF-17-020-01-CPPB) and National Institutes of Health (1T32HL134632-01 from the NHLBI) and indirectly by the National Cancer Institute (L30CA208950).

### Ethics Approval & Informed Consent

This study was reviewed and approved by the VA San Diego Healthcare System Institutional Review Board. All procedures performed in studies involving human participants were in accordance with the ethical standards of the institutional and/or national research committee and with the 1964 Helsinki Declaration and its later amendments or comparable ethical standards. Informed consent was obtained from all individual participants included in this study.

## Abbreviations

6MWD: = six-minute walk distance;
ANOVA: = one-way analysis of variance;
CI: = confidence interval;
COPD: = chronic obstructive pulmonary disease;
DL_CO_: = diffusion capacity of the lung for carbon monoxide;
EC: = exercise capacity;
LC13: = European Organization for Research and Treatment of Cancer QoL Questionnaire Core 30/Lung Cancer Module 13;
FEV1: = forced expiratory volume in 1 second;
HADS: = Hospital Anxiety and Depression Scale;
HF: = heart failure;
HFrEF: = heart failure with reduced ejection fraction;
MVA: = multivariable linear regression analysis;
NSCLC: = non-small cell lung cancer;
OR: = odds ratio;
PRO: = patient-reported outcome;
PTSD: = post-traumatic stress disorder;
QoL: = quality of life;
SE: = standard error;
SD: = standard deviation;
TLC: = total lung capacity;
SOBQ: = University California San Diego Shortness of Breath Questionnaire;
UVA: = univariable linear regression analysis;
VASDHS: = VA San Diego Healthcare System

