## Supplement Data for "Characterization of Dyspnea in Lung Cancer Survivors Following Curative-Intent Therapy"

**Supplemental Data**

E-Figure 1: Postulated Downward Spiral of Health in Lung Cancer Survivors Following Curative-Intent Therapy

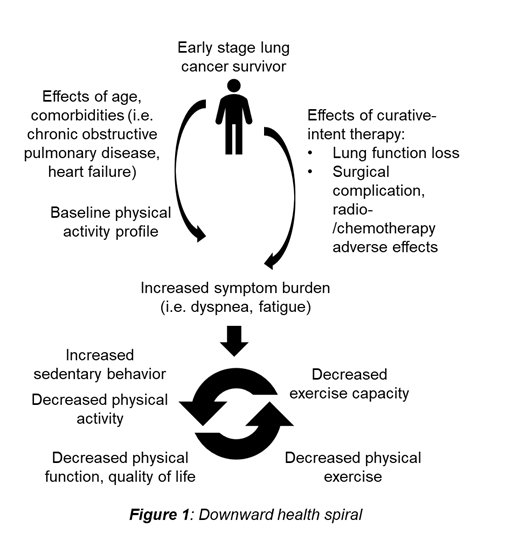

Description: Following curative-intent therapy, lung cancer survivors experience worsening symptom burden (i.e. dyspnea and fatigue), which contributes to more sedentary behavior and decreased physical activity. This decrease in physical activity leads to a progressive decline in physical function, which in turn decreases physical exercise behavior and exercise capacity. Over time, patients experience a downward spiral of health which goes unrecognized and negatively impacts quality of life. Interventions aimed at disrupting this vicious downward spiral may need to focus on **promptly** reducing symptom burden and/or encouraging sustainable physical activity to effectively improve physical function and quality of life in lung cancer survivors.

E-Figure 2: Scatter Plot of Dyspnea Scores

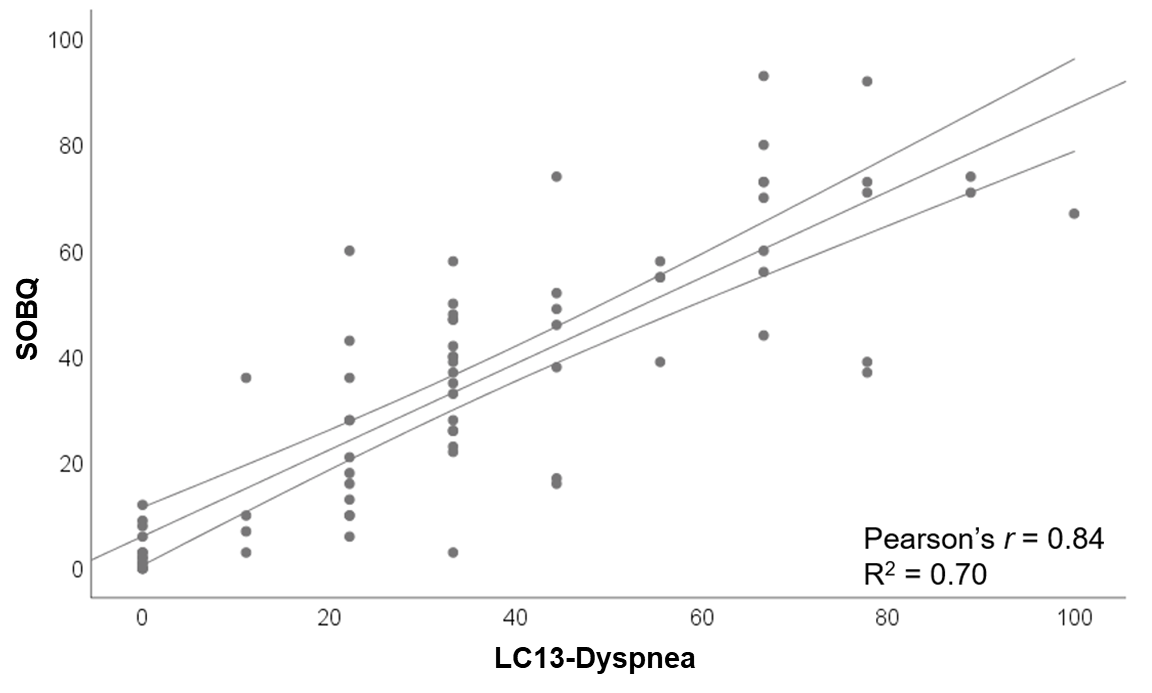

LC13 = European Organization for Research and Treatment of Cancer QoL Questionnaire Lung Cancer Module 13; SOBQ = University of California San Diego Shortness of Breath Questionnaire

E-Table 1: UVA – Significant/Borderline Determinants of Dyspnea

| *LC13-Dyspnea* | | | |
| --- | --- | --- | --- |
| Variable | β | R^2^ | P value |
| Age, year | -0.69 | 0.05 | 0.06 |
| HFrEF (N/Y) | -15.3 | 0.06 | 0.03 |
| Anxiety/depression/PTSD (N/Y) | -19.4 | 0.12 | 0.003 |
| FEV_1_ % predicted | -0.31 | 0.09 | 0.01 |
| DL_CO_  % predicted  Diffusion limited (N/Y) | -0.29  -10.7 | 0.08  0.04 | 0.02  0.08 |
| Lung hyperinflation (N/Y) | -15.5 | 0.07 | 0.04 |
| Primary treatment group | N/A | 0.07 | 0.08 |
| *SOBQ* | | | |
| Age, year | -0.63 | 0.04 | 0.08 |
| HFrEF (N/Y) | -16.5 | 0.08 | 0.02 |
| Anxiety/depression/PTSD | -19.9 | 0.13 | 0.002 |
| FEV_1_ % predicted | -0.37 | 0.13 | 0.002 |
| DL_CO_  % predicted  Diffusion limited (N/Y) | -0.33  -11.5 | 0.11  0.05 | 0.004  0.052 |
| Hyperinflation (N/Y) | -19.8 | 0.13 | 0.003 |
| Primary treatment group | N/A | 0.08 | 0.04 |

*^*^Partial R^2^*

DL_CO_ = diffusion capacity of the lung for carbon monoxide; LC13 = European Organization for Research and Treatment of Cancer QoL Questionnaire Lung Cancer Module 13; FEV_1_ = forced expiratory volume in 1 second; HFrEF = heart failure with reduced ejection fraction; PTSD = post-traumatic stress disorder; SOBQ = University of California San Diego Shortness of Breath Questionnaire

E-Table 2: UVA – Significant/Borderline Predictors of Functional EC

| Variable | β | R^2^ | P value |
| --- | --- | --- | --- |
| Age | -4.58 | 0.10 | 0.01 |
| Hyperlipidemia (N/Y) | 86.2 | 0.08 | 0.02 |
| CAD (N/Y) | 68.5 | 0.07 | 0.02 |
| FEV_1_ % predicted | 0.96 | 0.04 | 0.097 |
| DL_CO_ % predicted | 2.07 | 0.18 | <0.001 |
| Adenocarcinoma histology (N/Y) | -82.0 | 0.11 | 0.004 |
| Primary treatment group | N/A | 0.18 | 0.001 |
| Dyspnea scores, each point  LC13  SOBQ | -1.49  -1.85 | 0.10  0.15 | 0.01  0.001 |

β = regression coefficient; CAD = coronary artery disease; DL_CO_ = diffusion capacity of the lung for carbon monoxide; EC = exercise capacity; LC13 = European Organization for Research and Treatment of Cancer QoL Questionnaire Lung Cancer Module 13; FEV_1_ = forced expiratory volume in 1 second; SOBQ = University of California San Diego Shortness of Breath Questionnaire; UVA = univariable linear regression analysis
